# Comparing PAS domain coupled intrinsic dynamics in bHLH PAS domain transcription factor complexes

**DOI:** 10.1101/2025.05.07.652577

**Authors:** Karthik Sudarsanam, Ashutosh Srivastava, Sandhya P. Tiwari

## Abstract

The basic Helix-Loop-Helix–Per-Arnt-Sim (bHLH-PAS) transcription factors (TFs) are regulators of several critical cellular functions such as circadian rhythm, hypoxia response and neuronal development. These proteins contain tandemly repeated PAS domains that mediate heterodimer formation. While PAS domains adopt a conserved fold, recent studies suggest that their interaction interfaces differ distinctly in different TF complexes. However, the implications of these differences on the intrinsic dynamics of PAS domains remain unclear. In this study, we performed a comparative analysis of PAS domain dynamics across multiple bHLH-PAS TF complexes using all-atom Elastic Network Models (ENMs) and molecular dynamics (MD) simulations. We decomposed the intrinsic dynamics of PAS domains into self-coupled (internal domain dynamics) and directly coupled (interaction partner-influenced dynamics) motions using a projection-based approach. Our results show that self-coupled motions are more conserved across PAS domains than structure or sequence alone, while directly coupled motions capture the context-specific influence of partner proteins. Furthermore, hierarchical clustering of the overall covariance-based similarity scores revealed distinct grouping of CLOCK:BMAL1-type and HIF:ARNT-type complexes, which were not captured by sequence or structural comparisons. Root mean square fluctuation profiles derived from both MD and ENM approaches showed strong correspondence, validating the utility of ENMs in capturing biologically relevant dynamics, even in cases where the structural complexes were modelled using AlphaFold3. PAS-B domains were generally found to be less flexible than PAS-A domains for all the complexes analysed. Regions with high directly coupled flexibility were generally localized regions with high interface propensity in class I PAS-B domains, suggesting a higher level of coupled dynamics between PAS-B domains. Our results highlight how PAS domain intrinsic dynamics are shaped by both their internal architecture and complex-specific interactions, offering new insights into the functional diversification of bHLH-PAS transcription factors.

**Statement of Significance:** PAS domains are ubiquitous across all domains of life with diverse functions attributed to them. As part of bHLH-PAS transcription factors (TFs), they enable dimerization of Class I and Class II TFs. In this work, we investigated the effect of dimerization of PAS domains on their flexibility by using a method that allows us to isolate the intrinsic dynamics internal to a target domain and the intrinsic dynamics linked to the crosstalk between domains, from the all-atom elastic network model-based normal modes of the whole TF complex. Our findings reveal a context specific conservation of intrinsic dynamics based on the type of heterodimer complex. We also find a strong agreement between more-detailed MD simulations and the coarse-grained method used.

## Introduction

Per-Arnt-Sim (PAS) domains are ubiquitously found across the tree of life. They serve diverse functions in diverse groups of organisms, primarily acting as sensors (1,2). These primarily cytoplasmic sensor domains have evolved to recognise a wide variety of small molecules such as Heme and Flavins (3). Apart from sensing, PAS domain-containing proteins are known to play crucial role in several other cellular functions such as immune response (4), circadian rhythms (5), hypoxia response (6), etc. In many PAS domain-containing proteins, the function is also dependent on interactions with other structural and functional domains. Consequently, PAS domains occur in a variety of domain architectures. One of the major classes of PAS domain-containing proteins is the basic Helix Loop Helix-PAS domain transcription factors (TFs). These TFs, found in most metazoans, contain either one or two PAS domains (tandem repeats) along with a DNA binding domain. The bHLH-PAS domain TFs play roles in circadian rhythms, hypoxia response, neuronal development, etc. bHLH-PAS TFs function mostly as heterodimers with PAS domains playing critical role in the dimerization (7). Although some cofactor or ligand binding has been attributed to PAS domains in these TFs (8), conclusive evidence for many is still lacking. Thus, in terms of function, PAS domains in bHLH-PAS TFs primarily engage in the formation of protein-protein interaction, contributing to complex formation.

In case of mammals, there are primarily two classes of bHLH-PAS TFs (Class I and Class II) that form heterodimers to regulate the transcription of specific genes involved in various functions (9). Class I TFs with twelve members heterodimerize with Class II TFs containing four members (Table 1). In a recent analysis of PAS domain proteins from humans, it was reported that despite considerable sequence divergence, the PAS domain structures show remarkable similarity with canonical structural regions defined by four α-helices and five antiparallel β strands (10). The dimer structure conformations of the PAS domains were systematically analysed in the available crystal structures. As evident from the X-ray crystallography structures, despite considerable structural similarity, the PAS-A and PAS-B domains from Class I and Class II form very different interfaces when in complex. In CLOCK: BMAL1 complex, the PAS-A domains undergo a dimerization via the short α helix towards the N-terminal of the PAS domain, whereas the PAS-B domains use a different interface primarily occluding the β sheet of BMAL1 and the D, E and F α helices of CLOCK. Apart from this, the remaining protein-protein interfaces that are formed between PAS domains both within and across the chains are similar. The assembly of the bHLH-PAS TF heterodimer is an important step in complex formation. Despite several studies and reviews that have analysed various structural aspects, the dynamic implications of such different dimeric interfaces in PAS domains have not been studied extensively (3,7,9–14).

The dynamics of PAS domains from different proteins have been studied and compared using molecular dynamics (MD) simulations and subsequent extraction of essential dynamics (11,13). In one study, the authors performed short MD simulations, which revealed conservation in the dynamics in the β sheet region across different PAS domains (13). In another study, extensive structural modeling and molecular dynamics simulations were performed to find the dimerization interface between AhR and ARNT proteins (12). However, a comprehensive comparative analysis of the effects of dimerization on the dynamics of PAS domains from several different human bHLTH-PAS TF complexes has not been done, which we address in this study.

Here, we utilize all-atom elastic network models to capture the domain-domain coupled dynamics within a large set of bHLH-PAS TF complexes. First formulated by Tirion (15), elastic network models (ENMs) have allowed quick and computationally inexpensive normal mode analysis (NMA) that provide insight into the slow timescale dynamics of proteins. Since then, there have been many formulations of ENMs (16,17), and many applications including studies that examine the conservation of dynamics in protein superfamilies and structural folds(16,18–20). Furthermore, we use an implementation that extracts the intrinsic dynamics of a target domain (referred to as self-coupled motion) with explicit modelling of the full complex structure, and the intrinsic dynamics related to its crosstalk with a partner component (referred to as directly coupled motion) (21). This method has been shown to be effective in examining the promiscuous binding of ubiquitin with a variety of partner proteins (22,23) and involves the projection of submatrices from which the normal modes are reconstructed via singular value decomposition. The application of the coupled motions analysis method was modified to examine the coupling of the PAS domain to the rest of the transcription factor dimer complex, allowing us to dissect different aspects of protein-protein interactions on intrinsic dynamics.

We compared the intrinsic dynamics within target PAS domains (self-coupled) and the intrinsic dynamics influenced by partner protein-binding (directly coupled), extracted from the normal modes computed from the ENM of the full complex for each structure in the dataset. Furthermore, we corroborated with observations and found good agreement between the self-coupled intrinsic dynamics from MD simulations and ENM-based NMA. Our results indicate that the self-coupled intrinsic dynamics is more conserved across all PAS domains than structure and sequence. The directly coupled motions show a more nuanced picture of the influence of the partnering PAS domains, which is dependent on the complex context.

## Results and Discussion

### 1. Structure–sequence relationship of bHLH-PAS transcription factors

To understand the dynamic coupling between the PAS dimerization domains in the bHLH-PAS TFs we considered eleven members of the Class I TFs and two members of Class II TFs (Fig.1A, Table S1). All the transcription factors considered in this study share a conserved domain architecture, consisting of a bHLH DNA-binding domain followed by PAS-A and PAS-B domains. The PAS-A and PAS-B domains across all the TFs are similar in length (ranges 141-184 and 87-131 respectively). We first examined the sequence and structure-based relationships between all the PAS domains across the thirteen TFs considered here. We extracted the domain boundaries corresponding to the PAS-A and PAS-B domains for all the TFs and used the sequence corresponding to them and clustered them based on sequence similarity (Fig. S1A, Table S1). All the PAS-A domains irrespective of the Class of TFs, cluster together and similarly all the PAS-B domains cluster together, highlighting their evolutionary divergence (Fig. S1A). To further gain insights regarding the structure of PAS domains, we first modelled all the physiologically known complexes of human bHLH PAS TFs (Table S1). The PAS domain structures from each of the TFs were extracted, aligned and clustered based on root mean square deviation (RMSD). Here again, all the PAS-A and PAS-B domains cluster together respectively, following sequence similarity (Fig S1B). The PAS-A domain of NPAS4 shows higher divergence from other PAS-A domains, in structure-based clustering but not in sequence-based clustering. Despite these differences, the secondary structure elements are quite conserved across all the TFs following the canonical PAS domain topology (10) (Fig 1B). The differences are observed primarily in the loop regions. The bHLH, PAS A and PASB sequences for all the TFs considered were aligned and maximum likelihood based phylogenetic analysis was performed. Phylogenetic analysis and pairwise sequence similarity revealed distinct clustering patterns, with NPAS2 and CLOCK forming a close subgroup, AHR clustering with NPAS4, SIM1 grouping with SIM2, and NPAS1 with NPAS3. The hypoxia-inducible factors (HIF1A, HIF2A, and HIF3A) formed a separate clade, while ARNT1 clustered with BMAL1, reflecting their functional and evolutionary relationships (Fig. 1C).

**Figure 1.**
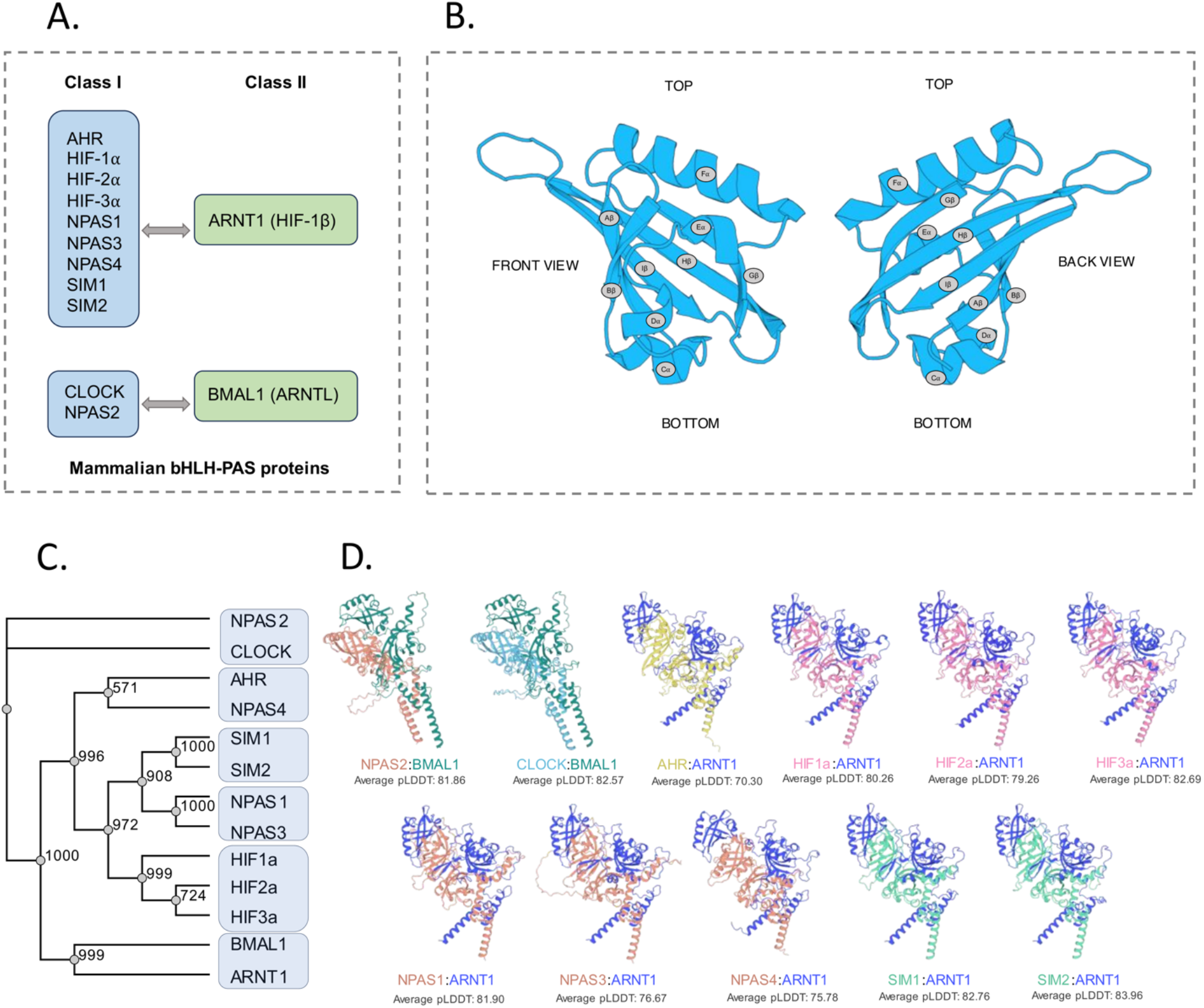
Sequence and structural survey of human bHLH-PAS transcription factors. (A) Class-I and class-II family of bHLH-PAS domain-containing proteins that are considered in this study (B) Structural representation of the PAS domain with highlighted secondary structure elements. (C) Phylogenetic tree illustrating the evolutionary relationships among bHLH-PAS transcription factors, with bootstrap values indicating the confidence of each branch. (D) Cartoon representation of the modeled complexes, shown in distinct colors, with corresponding average pLDDT values.

We then compared the structures of human bHLH PAS TFs. Considering that some of these TFs do not have experimentally derived structures, and there are missing residues in the ones that have known homologous X-ray crystallography structures, we modelled all the complex sequences using AlphaFold3 (AF3) (see methods). The predicted complexes exhibited high-confidence structural regions, with over 75% of pLDDT scores indicating reliable predictions (Fig. S1C). The pLDDT score was marginally better for the secondary structures as compared to the disordered and loop regions (Fig.S1C). We also compared the AF3 modelled structures with the experimentally resolved X-ray crystallography structures of the complexes and found very good correspondence between each pair.

RMSD based clustering of these complexes revealed two distinct clusters. (Fig. S1D). The comparisons of these complex structures further revealed that CLOCK: BMAL1 and NPAS2: BMAL1 adopt a similar conformation, distinct from other bHLH-PAS transcription factors in the family (Fig. 1D and S1D). Overall, there are two distinct heterodimer complex conformations that can be observed with considerable differences in the PAS-B domain orientation with respect to the rest of the complex (ROC). From here onwards we refer to these as CLOCK:BMAL1 type complex (or CB type complex) and HIF1α:ARNT1 type complex (or HA type complex) (Fig. S2). The PAS-A:PAS-A interface is similar across all heterodimers with the short N-terminal α helix (also refered to as A’ α helix) forming the interface with some small deviations. The PAS-B:PAS-B interface is also similar between the two distinct forms of the heterodimers (Fig. S2). The interface between PAS-B:PAS-B involves β sheet of Class II TF and the D, E and F α helices Class I TFs with residues (Fig. S2).

### 2. Covariance-based intrinsic dynamic coupling analysis in PAS domains captures protein flexibility in good agreement with MD simulations

To understand the effect of complex formation in Class I and Class II bHLH-PAS domain TFs on the dynamics of individual domains, we used all-atom ENM to extract normal modes and subsequent intrinsic dynamics. NMA generally captures the equilibrium motions of the system. To further validate the coarse-grained NMA results for the TF complexes, we first performed all-atom molecular dynamics simulations for one of the heterodimer complexes (CLOCK: BMAL1). The modelled structure of CLOCK: BMAL1 complex along with the cognate E-box (Enhancer-box) DNA was used as an initial model for performing all-atom MD simulations (Fig. 2A). One-microsecond MD simulation of this complex was performed in explicit solvent (see methods). The root mean square deviation (RMSD) analysis provided an overview of the overall structural deviations observed throughout the simulations, which stabilized after approximately 200 ns until the end of the simulation (Fig. 2B). The root mean square fluctuation (RMSF) analysis identified regions exhibiting higher flexibility, and the highest fluctuations were predominantly observed in the loop regions of both CLOCK as well as BMAL1 (Fig. S3A).

**Figure 2.**
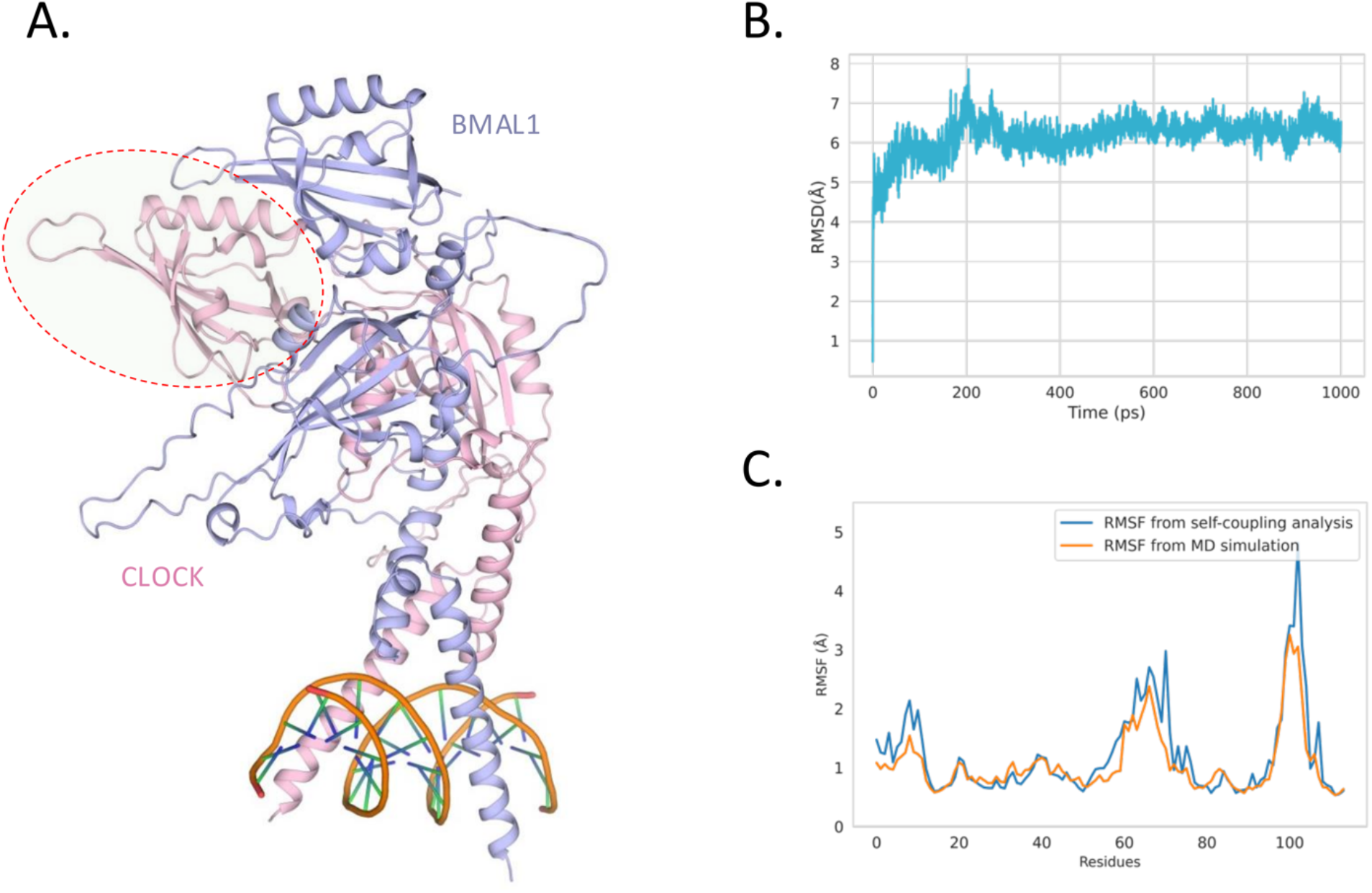
Molecular dynamics simulation results validate ENM self-coupled motions for CLOCK:BMAL1 complex. (A) Modeled *Homo sapiens* CLOCK:BMAL1:DNA (Enhancer-Box) complex used for simulation. The PAS-B domain of CLOCK is highlighted as circle. (B) Root Mean Square Deviation (RMSD) of the CLOCK:BMAL1 complex. (C) Line plot shows the comparison of Root Mean Square Fluctuation (RMSF) obtained from self-coupled analysis and Molecular Dynamics (MD) simulation for the PAS-B domain of CLOCK.

To compare the dynamics captured by NMA with the dynamics obtained through MD simulations, we focused on the PAS-B domain. The RMSF was calculated from MD simulations and by using the covariance matrix obtained from MD simulations. A good agreement could be observed between the RMSF obtained from these two methods highlighting the validity of using NMA to capture the intrinsic dynamics of these domains (Fig. 2C). We next investigated whether the correspondence in RMSF obtained via these two methods is dependent on the length of MD trajectories. For this, RMSF was calculated in two distinct ways. First, non-overlapping windows of 100 ns were used to calculate the covariance matrix followed by RMSF calculation (Fig S3B). We could observe a good correspondence among the RMSF calculated from different windows with some variations particularly in the first 100 ns trajectory results differing from the others. This could be attributed to the initial fluctuations in the complex structure at the start of dynamics. To further investigate the influence of each non-overlapping window on the overall RMSF profile, we calculated the RMSF from trajectory in cumulative time windows (0-100ns, 0-200 ns etc.). Here the results were much more uniform, and we could see that after 200 ns the subsequent windows gave very similar results, suggesting that the 200 ns dynamics was sufficient to capture the motions corresponding to the intrinsic dynamics of the complex from the covariance matrix (Fig. S3C).

### 3. Influence of binding partners on the intrinsic flexibility of PAS domains

Next, we analysed the intrinsic flexibility of PAS domains with explicit consideration of the rest of the TF complexes, by transforming the covariances obtained from all-atom NMA of the full complex into self-coupled and directly coupled motions using the method described by Dasgupta et al. (21–23). The coupled motions analysis was performed for PAS domains in Class I and Class II TFs in different complexes (see Methods). To determine the similarities in self-coupled and directly coupled motions, we pruned the covariance matrices to consist of only the structurally conserved regions in all the PAS domains, resulting in matrices of the same size. This was achieved by aligning all the PAS domain structures followed by sequence alignment and removing all the columns with any gaps in them, resulting in alignment of structurally equivalent residues across domains (Fig. S4A and B). The pruned covariances consisting of structurally conserved regions had dimensions of 88 × 88, and the Bhattacharya coefficient (BC) scores were calculated using bio3d, with additional hierarchical clustering of the BC scores (17,24).

We first compared the self-coupled motions in all PAS domains, essentially capturing the effect of the individual PAS domain’s internal motions extracted from the normal modes calculated for the full TF heterodimer complex. We observed that the BC scores for the self-coupled motions across all the PAS domains showed a high degree of similarity (Fig. 3A). Interestingly, PAS-A and PAS-B domains of BMAL1 in complex with CLOCK, show distinct dynamics as compared to all other PAS domains. Apart from this, there is a clear separation of PAS-A and PAS-B domains of TFs across all complexes, suggesting that the self-coupled motions are distinct in these two types of domains, even if they are found in the same TF complexes. Furthermore, we can observe that the PAS-A domains in the CB-type TF complexes form a separate cluster from the PAS-A domains of the HA-type TF complexes. Similarly, the PAS-B domains of the CB-type complexes form a separate subcluster from PAS-B domains of HA-type complexes. This shows that the method used is powerful enough to capture the encoded intrinsic dynamics information that distinguishes the domains from each context from submatrices that correspond to only the structurally conserved regions. We also observe that the derived self-coupled covariances capture the context of the two different types of complexes cleanly, which was not evident either in the sequence or structure-based clustering. Next, we analysed the effect of other domains and the cognate monomers on the intrinsic dynamics of each individual PAS domain. For this, the directly coupled motions were extracted from the full complex covariance matrices. Pairwise comparisons of the reconstructed directly coupled covariances revealed four major clusters (Fig. 3B). The ARNT PAS-B domain in HA-type complexes cluster together, and away from all other domains. Interestingly, the PAS-B domains of Class I TFs from CB-type complexes form a separate cluster from the PAS-B of Class I TFs from HA-type complexes, suggesting firstly, that the context of the complex is captured well in the direct coupling and secondly, that the effect on the intrinsic dynamics of PAS-B domain in both partnering TFs is asymmetric.

**Figure 3.**
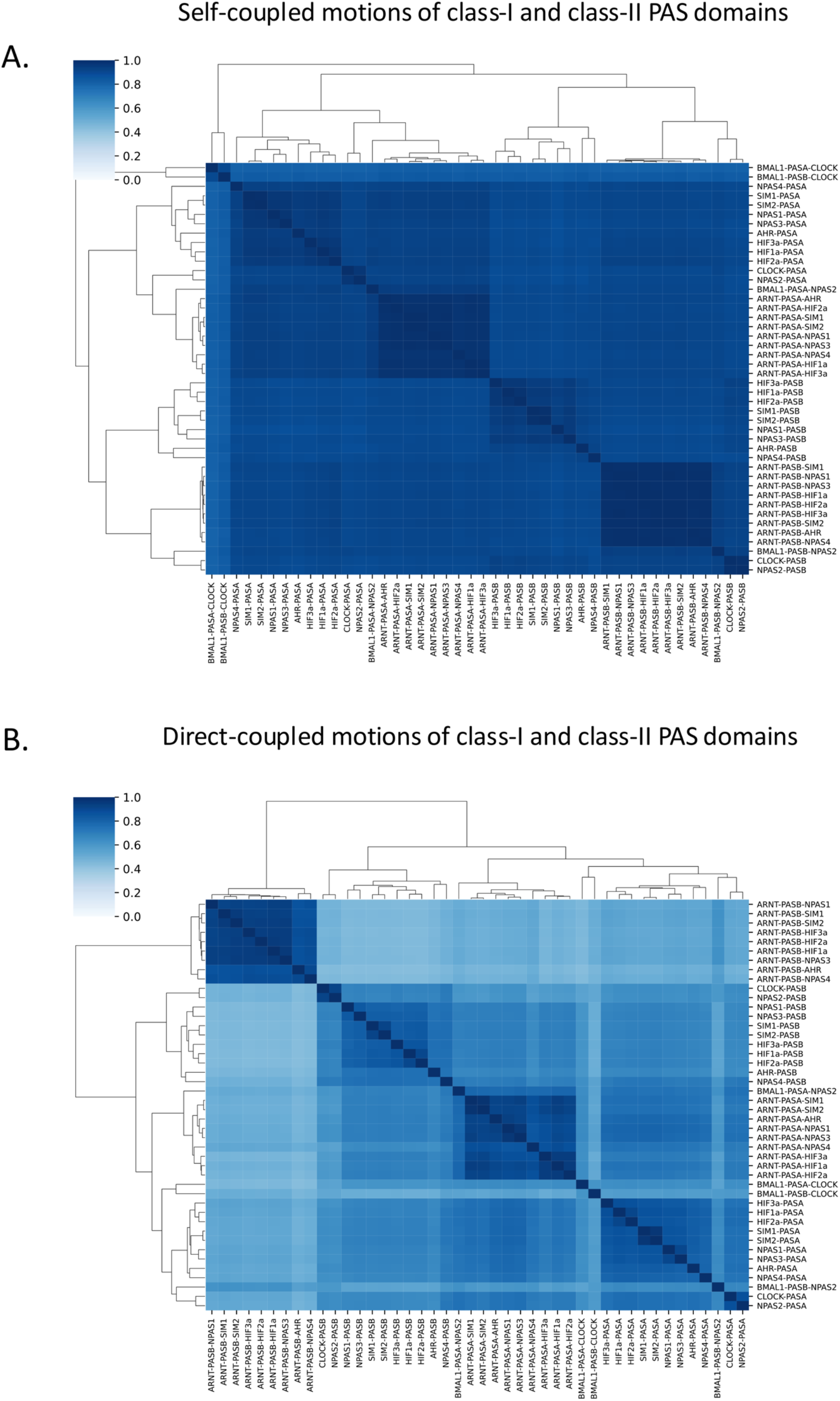
Heatmaps of self-coupled and directly coupled motions similarity. (A) Pairwise similarity comparison of covariance matrices from self-coupled analysis for PAS domains from Class-I and Class-II proteins is shown as heatmap. Each cell is colored according to the Bhattacharyya coefficient score. (B) Similarly, the Bhattacharyya coefficient score is applied to compare covariance matrices of Class-I and Class-II PAS domains from directly coupled analysis.

### 4. Correlation between structure, sequence, and intrinsic dynamics of PAS domains in bHLH-PAS transcription factors

Based on the previous analysis, we observed that the coupled motions analysis captures the context of complex formation better than the sequence or structure-based clustering. To further dissect the reason for this observation and to identify specific regions contributing to differences, we analysed the root mean square fluctuation (RMSF) derived from both self-coupled and directly coupled covariances. The RMSF profile of PAS-A from self-coupled motions showed a clear difference within the Class I TFs based on the type of complex (CB or HA) (Figure 4A). We further observe that within HA-type complexes, the NPAS4 PAS-A domain is an outlier. This is consistent with our earlier observation where NPAS4 PAS-A diverges from other Class I TFs based on structural clustering (Figure S1B). Similarly, PAS-A domain fluctuations in Class II TFs show clear differences based on the complex type (Figure 4B). Higher fluctuations can be observed in both Class I and II TFs in the loop between Gβ and Hβ, whereas only Class I TFs show higher fluctuations in the Fα helix region. The fluctuations here are higher in the CB-type complexes than in HA-type complexes. Similar observations were made for PAS-B domains where the RMSF shows divergence based on the complex type (Figure 4C and D). In general, much of the highest fluctuations were primarily observed in the loop regions, indicating that they are inherently flexible yet still dependent on the context of the ROC.

**Figure 4.**
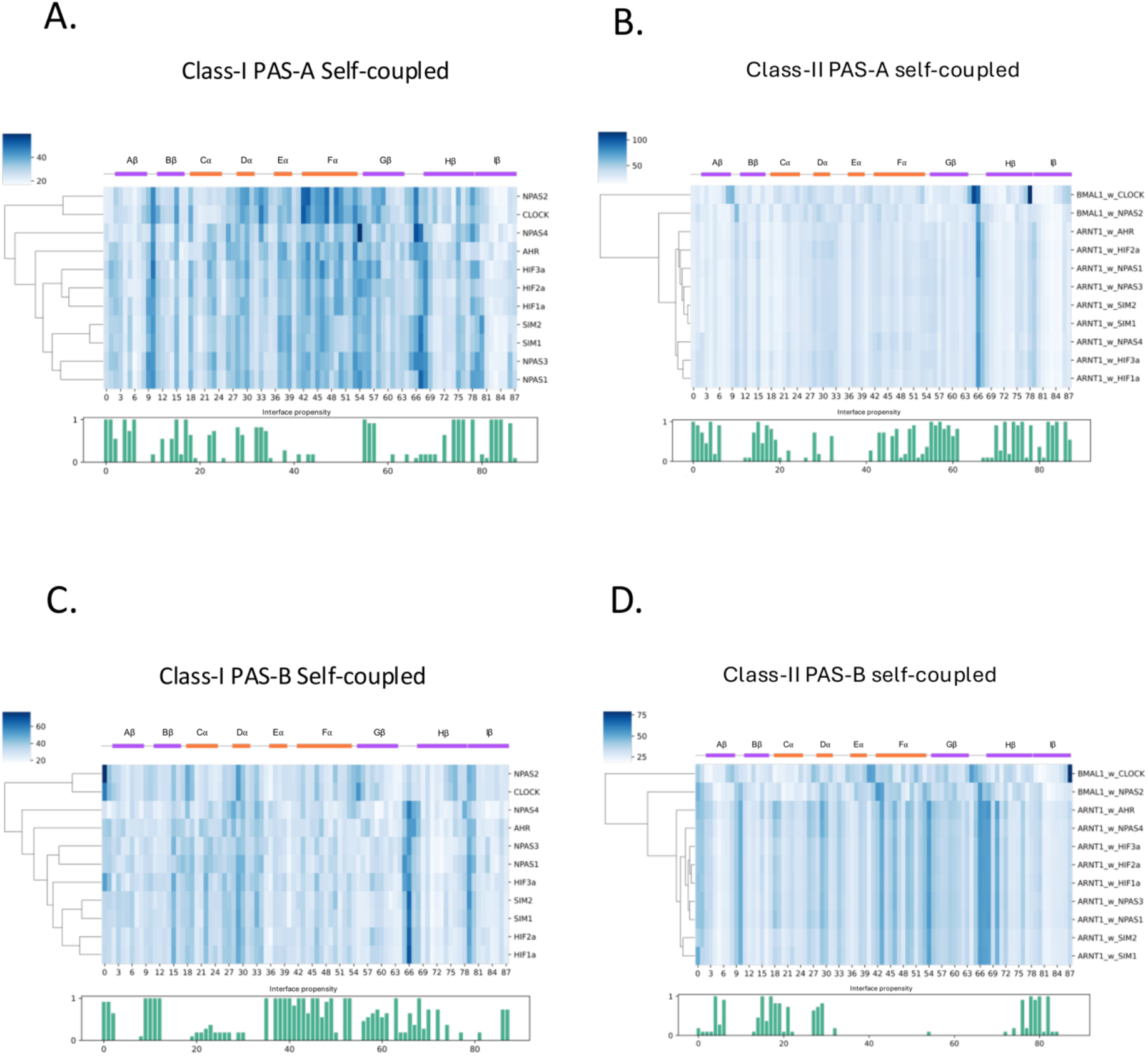
Self-coupled root mean square fluctuations (RMSF) and interface propensity profiles for class I and II PAS domains. (A-B) The residue-wise RMSF values of Class-I and Class-II PAS-A that are derived from self-coupled analysis are plotted as heatmap. Each column corresponds to the MSA index, whereas each row denotes a protein domain. The RMSF patterns of the PAS domains are hierarchically clustered. The secondary structure annotation is shown above the heat map. The interface propensity for each aligned position is shown as bar plot blow. (C-D) Similarly, residue-wise RMSF values for Class-I and Class-II PAS-B that are derived from self-coupled analysis are shown along with the interface propensity.

In the directly coupled motions analysis, which assesses the flexibility of individual PAS domains linked to the dynamic crosstalk with their binding partners, we observed distinct fluctuation patterns. We could also observe a similar pattern of hierarchical clustering based on fluctuations (Fig. S5). PAS-A in both Class I and II TFs, shows higher fluctuations in the Dα region but only in the HA-type complexes (Fig S5A and B). Within class II proteins, both PAS-A and PAS-B of ARNT1 displayed fluctuations primarily in the loop regions. However, in BMAL1 complexed with CLOCK and NPAS2, higher fluctuation patterns were observed in the Fα and Hβ strand. In the PAS-B domains of class I TFs, higher fluctuations were observed in Eα, Fα and Gβ region but not in class II TFs (Figure S5C and D). Specifically, the Cα helix showed increased flexibility in NPAS2 and CLOCK, while all proteins exhibited fluctuations in the Fα and Gβ regions (Fig. S5D).

To understand if the pattern of fluctuations observed originate in the dimer interface, we determined the interface propensity of the residues across the partnering domains (see methods). As expected, we observed that the RMSF from self-coupled motions does not correlate with interface propensity. Both class I and class II, PAS-A and PAS-B domains show an absence of high self-coupled RMSF at high interface propensity regions (Fig. 4). However, the RMSF from the directly coupled motions are relatively higher at regions with high interface propensity (Fig. 4 and S5). The Dα region shows consistently high directly coupled RMSF in class I PAS-A in all HA-type complexes. This region is at the interface between class I and class II PAS-A, suggesting strong coupling with cognate PAS-A domains. Similarly, in class I PAS-B domain, Eα, Fα regions with high interface propensity show consistently high directly coupled RMSF, whereas the Cα, Dα, and Hβ with residues of high interface propensities in class II TFs show even higher directly coupled RMSF than the rest of the sequence (Fig. S5 C and D). This suggests a higher degree of coupling between the PAS-B domains such that the influence of the ROC is manifested as higher directly coupled fluctuations at the dimer interface.

## Conclusions

PAS domains are ubiquitous and show considerable diversity in function, attracting significant interest in understanding their structure, evolution, and dynamics. Despite being primarily considered as sensors due to their ability to bind ligands and cofactors, the propensity of PAS domains to interact with other PAS domains plays a central role in many PAS domain-containing proteins. Particularly the bHLH-PAS transcription factors function as obligate heterodimers where the interaction between PAS domains is essential for the heterodimer formation. In this regard, understanding the structural, dynamics and evolutionary basis of this interaction can not only provide insights into the function but also in targeting them for therapeutic purposes. The sequence, structure and intrinsic dynamics analyses performed in this study have allowed us to investigate the extent to which the bHLH PAS domain transcription factors are conserved. Considering the high conservation in protein topology for PAS domains in general, we could pinpoint small but significant intrinsic dynamic differences at various levels - between i) PAS-A and PAS-B, ii) class I and class II, and iii) the sub-types within the TF complexes that we refer to as CB and HA-types. We demonstrated that the intrinsic dynamics within the PAS domains (self-coupled motions) are very similar across various complexes and that the subtle differences that we observe inform us about the state of the complex, hinting towards the distinct role of dynamics in complex formation. The directly coupled motions capture the effect of the rest of the complex on specific domains, showing a very clear pattern of influence according to complex type. We further show that despite very similar interfaces between the PAS domains, the intrinsic dynamics showed clear divergence in different types of complexes, which may not be evident from sequence or structure-based comparisons alone. The effectiveness of the coupled motion analysis in extracting both self-coupled and directly coupled fluctuations further highlighted key secondary structure regions of differences. In general, the fluctuations from self-coupled motions showed the regions that are affected by the motions of specific domains in the context of the rest of the complex. As expected, we observed low fluctuations due to self-coupled motions in the interface regions. In contrast, the directly coupled fluctuations showed a clear signal in terms of increased flexibility in the interface. However, even in the fluctuations, we could observe the divergence based on the types of complexes. We further demonstrated the robustness of elastic network models in reproducing root mean square fluctuations of the CLOCK: BMAL1 PAS-B domains from MD simulations and that with careful treatment of AlphaFold3 predicted models, we can augment missing structural information in large-scale studies. This work further paves the way for extracting the residue-specific contributions to the intrinsic dynamics by mining the correlation matrices and may further enable us to predict the potential interfaces between proteins that show promiscuous binding interactions with diverse interfaces.

## Methods

### Heterodimer structure modelling

The CLOCK: BMAL1 complex bound to DNA (Enhancer-Box), which was used for molecular dynamics simulations, was modelled by integrating AlphaFold2 multimer (25) predictions with the available crystal structure (PDB ID: 4H10). PAS-A and PAS-B domains were modelled using AlphaFold2, and the bHLH (DNA binding domain with E-box) domain from the X-ray crystallography structure (PDB ID: 4H10) was merged with this model to keep the protein DNA interactions intact. Energy minimization using GROMACS was performed as described later in the molecular dynamics simulation section.

The bHLH, PAS-A, and PAS-B domains of various bHLH-PAS domain-containing transcription factor complexes (Table S1) were modelled using AlphaFold v3 (26). The sequences corresponding to only these domains were extracted from the respective UniProt sequences and used for modelling. These complexes include AHR: ARNT1, CLOCK: BMAL1, HIF1α: ARNT1, HIF2α: ARNT1, HIF3α: ARNT1, NPAS1: ARNT1, NPAS2: BMAL1, NPAS3: ARNT1, NPAS4: ARNT1, SIM1: ARNT1, and SIM2: ARNT1. All the modelled complexes using AlphaFold v3 were energy minimized using UCSF Chimera v1.17.3 (27) with AMBER ff14SB force field with default parameters.

### Protein sequence and structure alignment

The sequences corresponding to PAS-A and PAS-B domains from all 13 TFs were extracted based on the domain boundaries as described in (Table S1). Multiple sequence alignment (MSA) was carried out using the CLUSTALW web server (28). Pairwise sequence similarity was calculated using Biopython (29). Hierarchical clustering was performed on the pairwise sequence similarity matrix using seaborn.clustermap (30).

For the phylogenetic tree construction, sequences corresponding to bHLH PAS-A and PAS-B domains for all 13 TFs were used as input for the MSA performed using the CLUSTALW web server (28). The MSA was used to construct a phylogenetic tree using Maximum likelihood (ML) method with 1000 bootstrap replicates in PhyML 3.0 (31).

PAS domain structures were extracted from all the complexes and aligned using MUSTANG v3.2.4 (32). The pairwise RMSD thus obtained was used for hierarchical clustering using in-house python (v3.10.13) script.

The structure-based sequence alignment obtained from MUSTANG (32) was pruned to remove any column with gaps, giving the final MSA comprising 88 columns of structurally equivalent residues across all PAS domains (Fig. S4A). This provided us with the consensus MSA which preserved the core structural alignment (Fig. S4B) while preventing inconsistencies in residue length.

The PAS-PAS domain interfaces were identified using MDAnalysis library (33) and an in-house python script. All the residues of Class-I TFs with any heavy atom within 5Å of class-II TFs and vice versa were treated as part of the interface in the complex. The interface residues of each PAS domain were mapped on the consensus MSA. The interface propensity was calculated as the ratio of number of interface residues in a column to the total number of residues in the column (11 for all cases). All the structures were visualised using PyMol v3.1.4 (34).

### Molecular Dynamics simulation

All-atom molecular dynamics (MD) simulations were performed for CLOCK: BMAL1 complex using GROMACS version 2021.3 (35). The Amber99sb-ildn force field was used, and the TIP3P water model was utilized to solvate the protein complexes within a dodecahedral simulation box. To maintain physiological ionic strength, Na+ and Cl-ions were added to maintain the concentration of 0.15 M. Energy minimization was performed using the steepest descent algorithm, with 50,000 steps or a maximum force of 500 kJ/mol/nm on any atom. Post-minimization, equilibration was carried out in two stages: first, 1 ns equilibration in the NVT ensemble, where the system temperature was maintained at 300 K using a modified Berendsen thermostat (36). This was followed by NPT equilibration in two sequential 1 ns steps, initially using the Berendsen barostat (36) and subsequently with the Parrinello-Rahman barostat (37), both at 1 bar pressure. Throughout, the system temperature was maintained at 300 K using the modified Berendsen thermostat (V-rescale). Finally, a production MD run of 1 microsecond was performed under the NPT ensemble using the Parrinello-Rahman barostat with pressure at 1 bar and temperature at 300 K. The root mean square deviation (RMSD), root mean square fluctuation (RMSF), covariance matrices were computed using in-built programs of GROMACS gmx rms, gmx rmsf and gmx covar.

### Normal Mode Analysis

All-atom elastic network model-based normal modes were calculated using the bio3d package (38,39) in R v4.0.0. In bio3d, the all-atom forcefield follows Hinsen et al 2000 (40), where the force constants were obtained by fitting to a local energy minimum of a crambin model derived from the AMBER99SB force field. Covariance matrices were calculated from the full computed normal modes, and the values for the Cα atom positions were subsequently used to calculate for coupled motions using in-house R scripts as implemented in Dasgupta et al. (21–23).

### Coupled Motions Analysis

Briefly, to derive self-coupled motions, the amino acid position index corresponding to the protein domain of interest was extracted from the multiple structural alignments, such that only fully conserved positions in the alignment were considered. Then, this amino acid position index formed the basis for projection to a subspace without the six external degrees of freedom associated with rigid-body rotations and translations between the domain of interest and the rest of the complex (ROC). Thus, the projected submatrix *P* that indexes only the domain of interest A (symmetrical) comprises covariances *C* due to its own intrinsic flexibility (referred to as self-coupled motion),

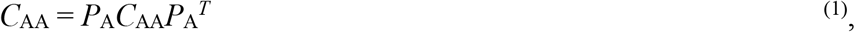

while the projected submatrix with the index of the domain of interest A and the rest of the structure B (asymmetrical) comprises covariances that correspond to the intrinsic flexibility influenced by the rest of the structure (referred to as directly coupled motion) as follows,

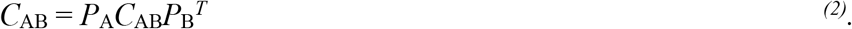

By performing singular value decomposition (SVD) on the projected covariance submatrices, we obtain self-coupled motions from each pair component of the structural complex and the directly coupled motions between those components. The reconstructed covariance matrices obtained from SVD eigenvalues have the same square dimensions of the larger sequence index and were then used to derive the root mean squared fluctuation profiles (related to the trace of the diagonal) for both the self-coupled and directly coupled motions.

### Clustering of covariance matrices

To compare the covariance matrices of PAS domains between class-I and class-II proteins, the Bhattacharyya coefficient (24), as implemented in bio3d, was used as a measure of similarity. The covariance matrices were first trimmed to 88×88 corresponding to the consensus MSA.

These covariances matrices converted into probability distributions, and the Bhattacharyya coefficient was then computed to quantify the overlap between these distributions. Additionally, hierarchical clustering was performed using the python package seaborn.clustermap (30) function to visualize the relationships between the covariance matrices.

## Data availability

All the raw data and custom scripts used in this study are publicly available on Zenodo at the following link:<u>10.5281/zenodo.15349957</u>.

## Author contributions

KS performed research, analyzed data and wrote manuscript. AS designed research, analyzed data, wrote and edited manuscript. SPT designed research, analyzed data, wrote and edited manuscript.

## Acknowledgements

KS acknowledges financial support from IIT Gandhinagar excellence fellowship. AS acknowledges financial support from DBT-Ramalingaswami Fellowship and IIT Gandhinagar for intramural funding. The PARAM Ananta Supercomputing facility is acknowledged for computational resources. This work was (in part) performed under the International Collaborative Research Program of Institute for Protein Research, Osaka University, ICR-24-16 to SPT.

## References

1. Gu YZ, Hogenesch JB, Bradfield CA. The PAS Superfamily: Sensors of Environmental and Developmental Signals. Annu Rev Pharmacol Toxicol. 2000 Apr;40(1):519–61.

2. Xing J, Gumerov VM, Zhulin IB. Origin and functional diversification of PAS domain, a ubiquitous intracellular sensor. Sci Adv. 2023 Sep;9(35):eadi4517.

3. Möglich A, Ayers RA, Moffat K. Structure and Signaling Mechanism of Per-ARNT-Sim Domains. Structure. 2009 Oct;17(10):1282–94.

4. Xue X, Ramakrishnan S, Anderson E, Taylor M, Zimmermann EM, Spence JR, et al. Endothelial PAS Domain Protein 1 Activates the Inflammatory Response in the Intestinal Epithelium to Promote Colitis in Mice. Gastroenterol.. 2013 Oct;145(4):831–41.

5. Gustafson CL, Partch CL. Emerging Models for the Molecular Basis of Mammalian Circadian Timing. Biochem. 2015 Jan 20;54(2):134–49.

6. Scheuermann TH, Yang J, Zhang L, Gardner KH, Bruick RK. Hypoxia-Inducible Factors Per/ARNT/Sim Domains: Structure and Function. Methods in Enzymol. 2007;435:3–24

7. Sharma D, Partch CL. PAS Dimerization at the Nexus of the Mammalian Circadian Clock. J Mol Biol. 2024 Feb;436(3):168341.

8. Freeman SL, Kwon H, Portolano N, Parkin G, Venkatraman Girija U, Basran J, et al. Heme binding to human CLOCK affects interactions with the E-box. Proc Natl Acad Sci. 2019 Oct;116(40):19911–6.

9. Daffern N, Radhakrishnan I. Per-ARNT-Sim (PAS) Domains in Basic Helix-Loop-Helix (bHLH)-PAS Transcription Factors and Coactivators: Structures and Mechanisms. J Mol Biol. 2024 Feb;436(3):168370.

10. Rojas BL, Vazquez-Rivera E, Partch CL, Bradfield CA. Dimerization Rules of Mammalian PAS Proteins. J Mol Biol. 2024 Feb;436(3):168406.

11. Vreede J, Van Der Horst MA, Hellingwerf KJ, Crielaard W, Van Aalten DMF. PAS Domains. J Biol Chem. 2003 May;278(20):18434–9.

12. Corrada D, Soshilov AA, Denison MS, Bonati L. Deciphering Dimerization Modes of PAS Domains: Computational and Experimental Analyses of the AhR:ARNT Complex Reveal New Insights Into the Mechanisms of AhR Transformation. PLOS Comput Biol. 2016 Jun 13;12(6):e1004981.

13. Pandini A, Bonati L. Conservation and specialization in PAS domain dynamics. Protein Eng Des Sel. 2005 Mar 1;18(3):127–37.

14. Dong Z, Zhou H, Tao P. Combining protein sequence, structure, and dynamics: A novel approach for functional evolution analysis of PAS domain superfamily. Protein Sci. 2018 Feb;27(2):421–30.

15. Tirion MM. Large Amplitude Elastic Motions in Proteins from a Single-Parameter, Atomic Analysis. Phys Rev Lett. 1996 Aug 26;77(9):1905–8.

16. Tiwari SP, Reuter N. Conservation of intrinsic dynamics in proteins — what have computational models taught us? Curr Opin Struct Biol. 2018 Jun;50:75–81.

17. Fuglebakk E, Tiwari SP, Reuter N. Comparing the intrinsic dynamics of multiple protein structures using elastic network models. Biochim Biophys Acta BBA - Gen Subj. 2015 May;1850(5):911–22.

18. Leo-Macias A, Lopez-Romero P, Lupyan D, Zerbino D, Ortiz AR. An Analysis of Core Deformations in Protein Superfamilies. Biophys J. 2005 Feb;88(2):1291–9.

19. Yang L, Song G, Carriquiry A, Jernigan RL. Close Correspondence between the Motions from Principal Component Analysis of Multiple HIV-1 Protease Structures and Elastic Network Modes. Structure. 2008 Feb;16(2):321–30.

20. Tiwari SP, Reuter N. Similarity in Shape Dictates Signature Intrinsic Dynamics Despite No Functional Conservation in TIM Barrel Enzymes. Jernigan RL, editor. PLOS Comput Biol. 2016 Mar 25;12(3):e1004834.

21. Dasgupta B, Tiwari SP. Explicit versus implicit consideration of binding partners in protein–protein complex to elucidate intrinsic dynamics. Biophys Rev. 2022 Dec;14(6):1379–92.

22. Dasgupta B, Nakamura H, Kinjo AR. Counterbalance of ligand- and self-coupled motions characterizes multispecificity of ubiquitin. Protein Sci. 2013 Feb;22(2):168–78.

23. Dasgupta B, Nakamura H, Kinjo AR. Rigid-body motions of interacting proteins dominate multispecific binding of ubiquitin in a shape-dependent manner. Proteins Struct Funct Bioinformatics. 2014 Jan;82(1):77–89.

24. Fuglebakk E, Echave J, Reuter N. Measuring and comparing structural fluctuation patterns in large protein datasets. Bioinformatics. 2012 Oct 1;28(19):2431–40.

25. Evans R, O’Neill M, Pritzel A, Antropova N, Senior A, Green T, et al. Protein complex prediction with AlphaFold-Multimer. bioRxiv; 2021; doi: 10.1101/2021.10.04.463034

26. Abramson J, Adler J, Dunger J, Evans R, Green T, Pritzel A, et al. Accurate structure prediction of biomolecular interactions with AlphaFold 3. Nature. 2024 Jun 13;630(8016):493–500.

27. Pettersen EF, Goddard TD, Huang CC, Couch GS, Greenblatt DM, Meng EC, et al. UCSF Chimera—A visualization system for exploratory research and analysis. J Comput Chem. 2004 Oct;25(13):1605–12.

28. Thompson JD, Higgins DG, Gibson TJ. CLUSTAL W: improving the sensitivity of progressive multiple sequence alignment through sequence weighting, position-specific gap penalties and weight matrix choice. Nucleic Acids Res. 1994;22(22):4673–80.

29. Cock PJA, Antao T, Chang JT, Chapman BA, Cox CJ, Dalke A, et al. Biopython: freely available Python tools for computational molecular biology and bioinformatics. Bioinformatics. 2009 Jun 1;25(11):1422–3.

30. Waskom M. seaborn: statistical data visualization. J Open Source Softw. 2021 Apr 6;6(60):3021.

31. Guindon S, Dufayard JF, Lefort V, Anisimova M, Hordijk W, Gascuel O. New Algorithms and Methods to Estimate Maximum-Likelihood Phylogenies: Assessing the Performance of PhyML 3.0. Syst Biol. 2010 Mar 29;59(3):307–21.

32. Konagurthu AS, Whisstock JC, Stuckey PJ, Lesk AM. MUSTANG: A multiple structural alignment algorithm. Proteins Struct Funct Bioinforma. 2006 Aug 15;64(3):559–74.

33. Gowers R, Linke M, Barnoud J, Reddy T, Melo M, Seyler S, et al. MDAnalysis: A Python Package for the Rapid Analysis of Molecular Dynamics Simulations. 2016; In S. Benthall and S. Rostrup, editors, Proceedings of the 15th Python in Science Conference, pages 98–105, Austin, TX

34. Janson G, Paiardini A. PyMod 3: a complete suite for structural bioinformatics in PyMOL. Arne E, editor. Bioinformatics. 2021 Jun 16;37(10):1471–2.

35. Páll S, Zhumarov A, Bauer P, Abraham M, Lundborg M, Gray A, et al. Heterogeneous parallelization and acceleration of molecular dynamics simulations in GROMACS. J. Chem. Phys. 2020 Oct; 153, 134110

36. Berendsen HJC, Postma JPM, Van Gunsteren WF, DiNola A, Haak JR. Molecular dynamics with coupling to an external bath. J Chem Phys. 1984 Oct 15;81(8):3684–90.

37. Parrinello M, Rahman A. Polymorphic transitions in single crystals: A new molecular dynamics method. J Appl Phys. 1981 Dec 1;52(12):7182–90.

38. Grant BJ, Rodrigues APC, ElSawy KM, McCammon JA, Caves LSD. Bio3d: an R package for the comparative analysis of protein structures. Bioinformatics. 2006 Nov 1;22(21):2695– 6.

39. Grant BJ, Skjærven L, Yao X. The BIO3D packages for structural bioinformatics. Protein Sci. 2021 Jan;30(1):20–30.

40. Hinsen K, Petrescu AJ, Dellerue S, Bellissent-Funel MC, Kneller GR. Harmonicity in slow protein dynamics. Chem Phys. 2000 Nov;261(1–2):25–37.

