## Supplementary File for "Comparing PAS domain coupled intrinsic dynamics in bHLH PAS domain transcription factor complexes"

**Supporting material**

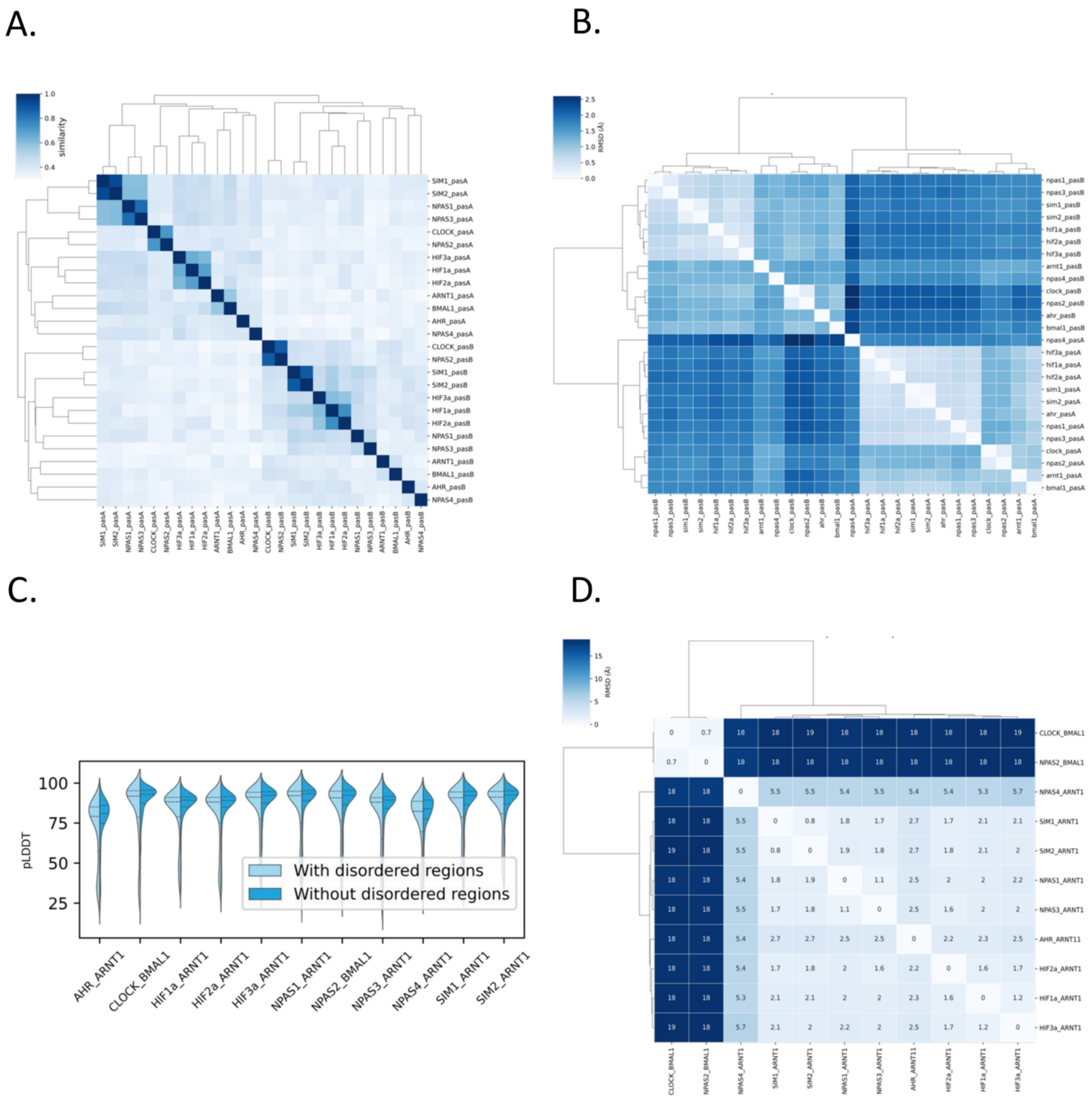


**Figure S1.** (A) Heatmap depicting the pairwise sequence similarity of PAS-A and PAS-B domains from Class-I and Class-II proteins, with hierarchical clustering. The similarity values are represented using a colour gradient, as indicated by the colour bar. (B) Similarly, the pairwise RMSD comparison of PAS-A and PAS-B domains in Class-I and Class-II proteins are shown. (C) pLDDT distribution of modelled bHLH-PAS domains with and without disordered regions. (D) Heatmap of pairwise RMSD comparisons for bHLH-PAS domain transcription factor complexes. The RMSD values were computed to assess structural similarities, with hierarchical clustering. Values are color-coded according to the colour bar.

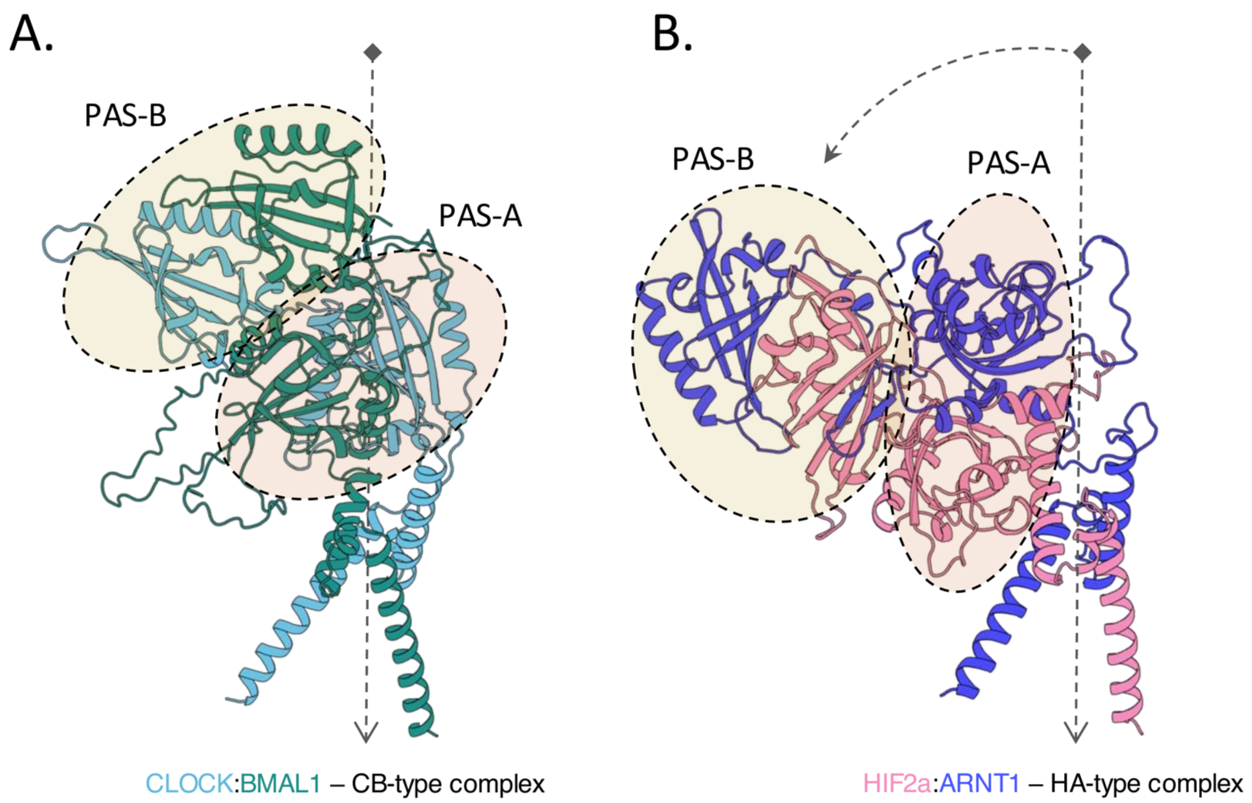


**Figure S2.** (A) To represent the CB-type complexes, the conformation of CLOCK:BMAL1 is shown with PAS-A and PAS-B domains highlighted. (B) Similarly, the conformation of HIF2a:ARNT1 is shown to represent the HA-type complexes. The arrow indicates the extent of conformational changes observed in type-HA compared to type-CB PAS domain.

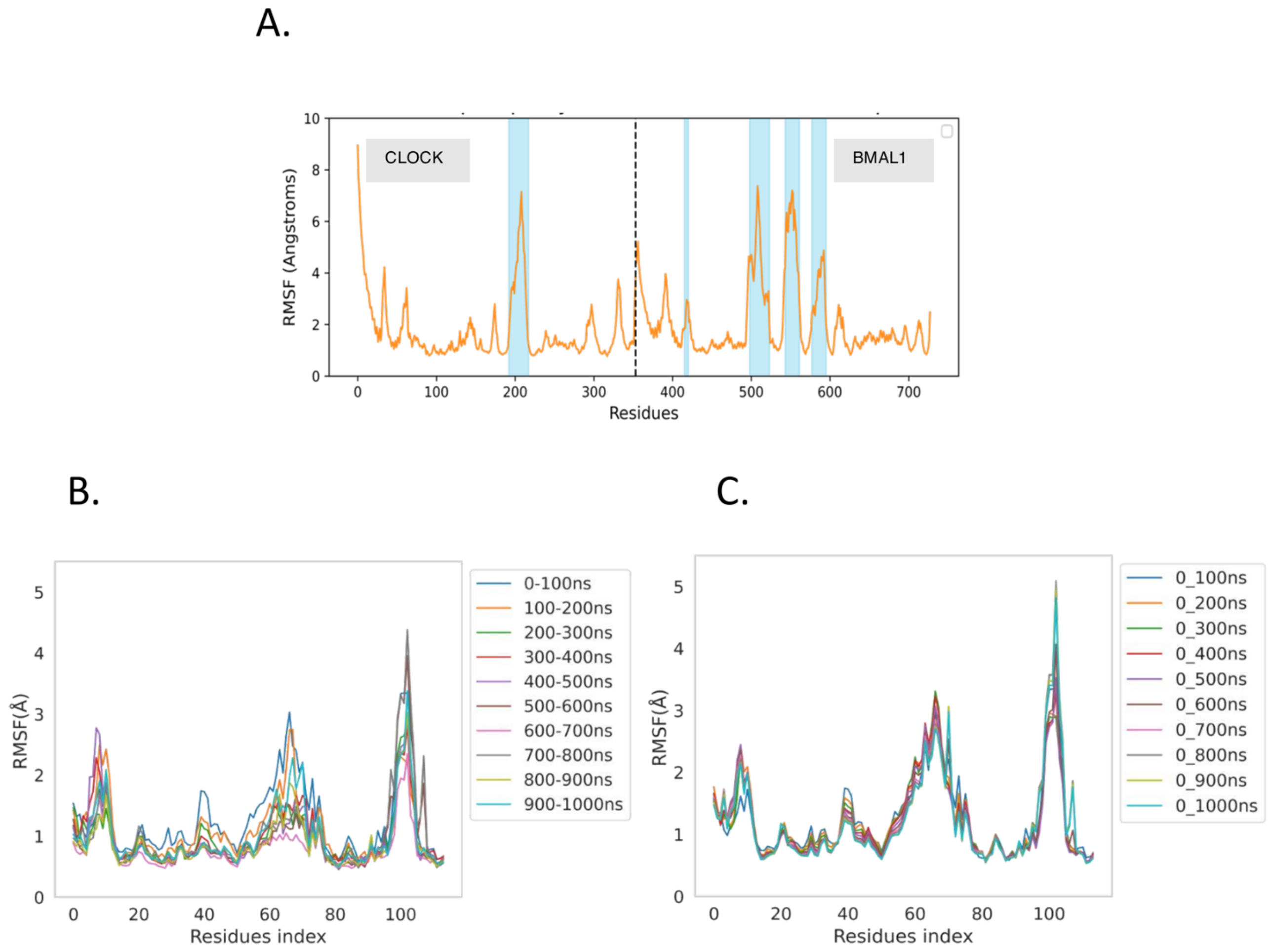


**Figure S3.** (A) RMSF of the CLOCK:BMAL1 complex from MD simulations, with disordered regions highlighted. (B-C) Root mean square fluctuation (RMSF) of the PAS-B domain of CLOCK obtained from self-coupled analysis across different simulation windows.

A.

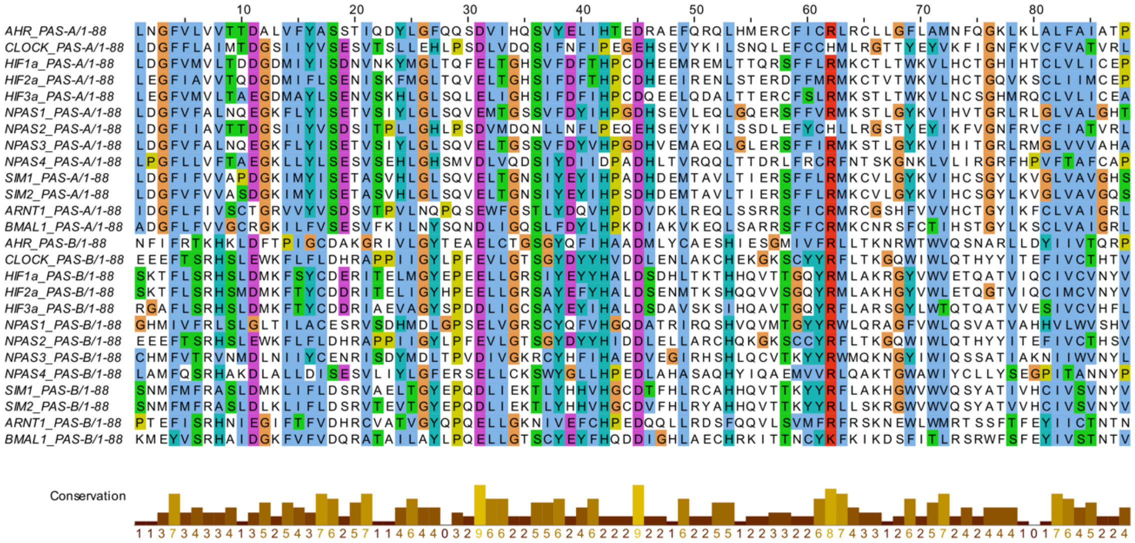


B.

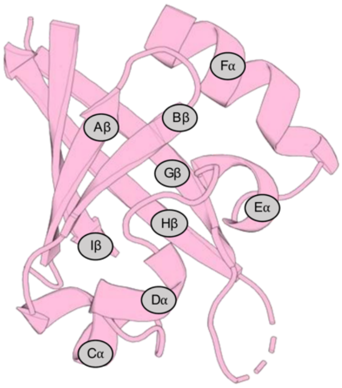


**Figure S4.** (A) Refined multiple sequence alignment of PAS domains from Class-I and Class-II proteins after gap removal, visualized using the Clustal-X color scheme. The conservation scores are shown below the multiple sequence alignment (MSA) as a bar graph, highlighting regions of high and low conservation. (B) Cartoon representation of the PAS domain illustrating the conserved core structure retained after gap removal from the structure-based sequence alignment.

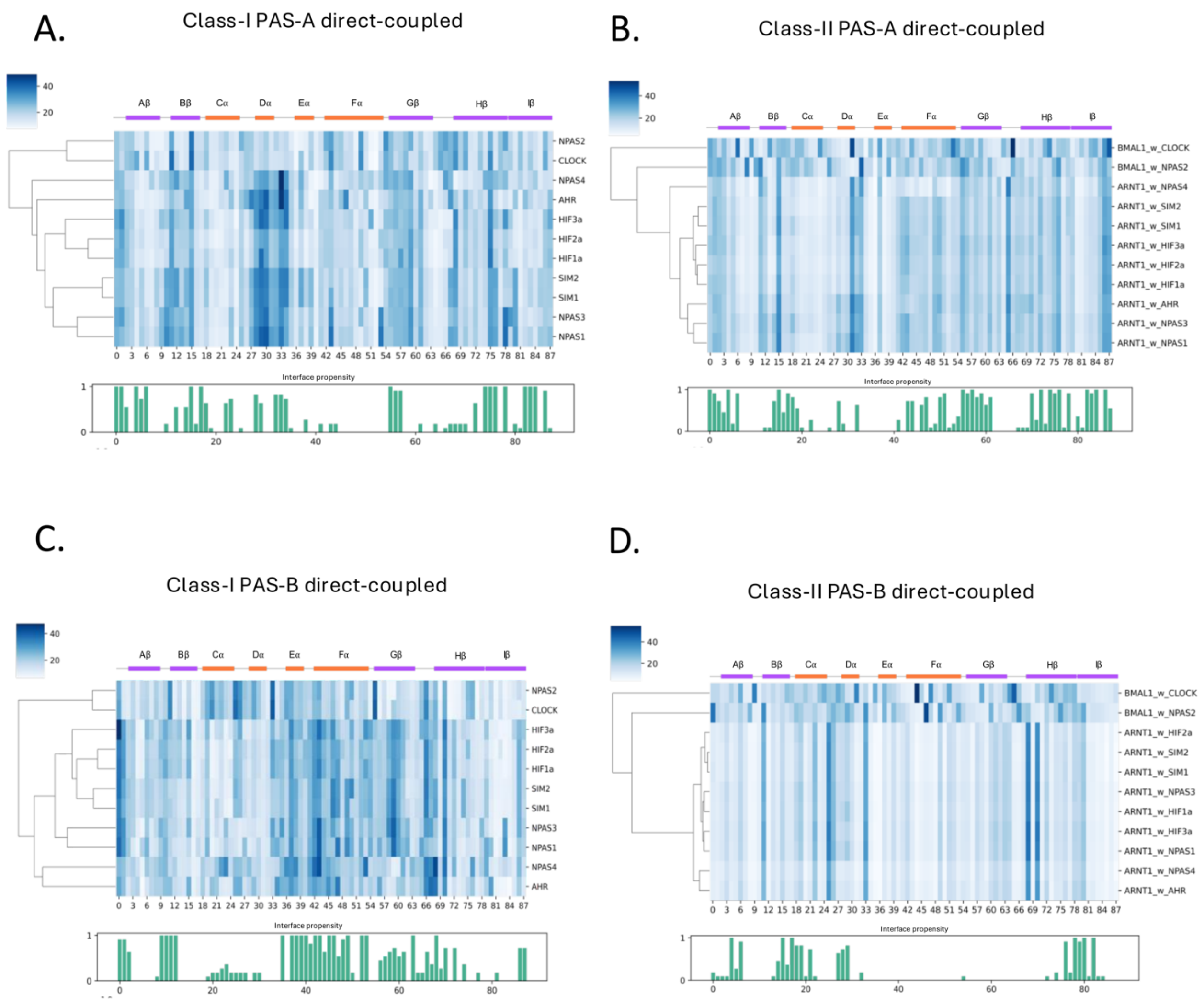


**Figure S5.** (A-B) Heatmap of residue-wise RMSF values for Class-I and Class-II PAS-A domains, derived from direct-coupling analysis. Each column represents the MSA index, while each row corresponds to a protein domain. Hierarchical clustering highlights RMSF patterns across PAS domains. Secondary structure annotations are shown above the heatmap. Below, interface propensity is represented as a bar plot. (C-D) Similarly, residue-wise RMSF values for Class-I and Class-II PAS-B that are derived from direct-coupled analysis are shown along with the interface propensity.

| No | Protein Names | UniProt ID (Homo sapiens) | BHLH residues | PAS-A residues | PAS-B residues |
|---|---|---|---|---|---|
| 1. | ARNT | P27540 | 89-142 | 161-235 | 349-467 |
| 3. | BMAL1 | O00327 | 71-125 | 143-215 | 326-444 |
| 4. | NPAS1 | Q99742 | 45-98 | 135-207 | 293-408 |
| 5. | NPAS3 | Q8IXF0 | 51-104 | 147-217 | 319-406 |
| 6. | NPAS4 | Q8IUM7 | 1-53 | 70-144 | 203-317 |
| 7. | HIF1a | Q16665 | 17-70 | 85-158 | 228-345 |
| 8. | HIF2a | Q99814 | 14-67 | 84-154 | 230-347 |
| 10. | HIF3a | Q9Y2N7 | 14-67 | 82-154 | 227-358 |
| 11. | AHR | P35869 | 27-80 | 111-181 | 275-386 |
| 14. | CLOCK | O15516 | 34-84 | 107-177 | 262-379 |
| 13. | SIM1* | P81133 | 1-53 | 77-147 | 218-335 |
| 14. | SIM2* | Q14190 | 1-53 | 77-149 | 218-335 |
| 15. | NPAS2* | Q99743 | 9-59 | 82-152 | 237-354 |

* *Experimental Structures are not available*

**Table S1.** Sequence ranges of bHLH, PAS-A, and PAS-B domains in Class-I and Class-II transcription factors (TFs).
